# Lumbar time-varying muscle synergies in trunk flexion and bending movements at different velocities

**DOI:** 10.1101/2022.04.03.486861

**Authors:** Mahdi Bagheri Rouchi, Mehrdad Davoudi, Narges Meftahi, Ehsan Rashedi, Mohamad Parnianpour, Kinda Khalaf

## Abstract

**Purpose:** Although the extent of which the central nervous system uses muscle synergies as a movement control strategy remains an open area of research, it is widely agreed that synergies facilitate the robustness of the neuromuscular system, allowing for effective postural control and flexible movement. This work aimed to investigate the muscle activation patterns of the trunk and time-varying muscle synergies using a novel 18-muscle 3-DOF, 3-D musculoskeletal model of the lumbar spine developed by the authors.

**Methods:** 24 different biaxial trunk movements were simulated via the optimization of kinetic and kinematic measures towards obtaining the corresponding muscle activation patterns at 3 different velocities. These patterns were subsequently used to extract the principal (phasic and tonic) spatio-temporal synergies associated with the observed muscle activation patterns in the range of simulated movements.

**Results:** Four dominant synergies were able to explain a considerable percent (about 75%) of the variance of the simulated muscle activities. The extracted synergies were spatially tuned in the direction of the main simulated movements (flexion/extension and right/left lateral bending). The temporal patterns demonstrated gradual monotonic shifts in tonic synergies and biphasic modulatory components in phasic synergies with spatially tuned time-delays. The increase in velocity resulted in an elevated amplitude coefficient and accelerated activation of phasic synergies.

**Conclusion:** Our results suggest the plausibility of a time-varying synergies strategy in the dynamic control of trunk movement. Further work is needed to explore leveraging these concepts in various applications, such as rehabilitation and musculoskeletal biomechanics, towards providing more insight into the mechanisms underlying trunk stability and flexibility.

## Introduction

The sophisticated design of the lumbar spine, integrating a large number of muscles and high degrees of freedom, allows for the inherent redundancy, also referred to as functional abundance, to be well leveraged by the trunk’s motor control, especially during complex combined movements (1-3). Among the variety of proposed modular control strategies on both the kinematic level (kinematic modularity) (4, 5), and execution level (execution modularity) (6, 7), the muscle synergies control strategy provides a plausible solution to the “ill-posed” control problem. A muscle synergy is in essence a functional unit coordinating the activity of several muscles, such that the activation of a group of muscles contributes to a particular movement, hence reducing the dimensionality of muscle control (8-10). Previous experimental findings have supported the principle of synergistic control of abdominal and back muscles in the trunk region during isometric lumbar exertion (8, 11). On the other hand, is still debatable whether such experimental observations reflect a physiological basis of low-dimensional control in the central nervous system (CNS), or an epiphenomenon based on task constraints and/or biomechanics (12).

Muscle synergies extraction has been performed by various decomposition algorithms based on two general approaches: synchronous synergies and time-varying synergies (13, 14). Based on synchronous synergies, synergistic groups of muscles are activated synchronously with a fixed time course. One of the main limitations of this approach is the lack of time delay between the onset of muscle activation for each synergy, despite evidence that in complex movements, muscles are often activated asynchronously (10, 15). The onset time of muscle activity included in many movements may relatively change with different movement directions and velocities (16, 17), hence rendering the synchronous synergies theory too simplistic for the interpretation of coordinated muscles activities (15). The time-varying synergies model addresses this limitation by incorporating both the spatial characteristics of muscle activation, as well as the temporal features (10, 18). According to this control scheme, basic units of motor production not only involve the fixed coordination of relative muscle activation amplitudes, but also the coordination of activation timings. Time-varying synergies generally forms sequence units, where temporally coordinated (but not necessarily synchronous) drives are supplied to groups of muscles (19). d’Avella et al. included such time shifts in their optimization model, in addition to amplitude coefficients, to introduce time-varying synergies for better explanation of the behavior of the CNS in controlling the shoulder and hand in fast-reaching movements (7). They showed that while both time-varying and constant synergies refer to a common concept of CNS function, time-varying synergies were able to provide a better explanation of the original dataset with a smaller number of synergies (7).

To the best of our knowledge, no previous study has evaluated the impact of different velocities and planes of motion during trunk flexion-extension movement on amplitude coefficients and the onset time of synergies patterns. Using an optimization algorithm, we have extracted spatially tuned time-varying muscle synergies for lumbar flexural movements. These synergies included proper time scales of delays to reconstruct movements with multiple velocities. Because of the limitations of superficial electromyography (e.g., lack of access to deep muscles), we have developed an 18-muscle 3-D model of the human lumbar spine to estimate muscular activity. Furthermore, it has been reported that separation of muscle activation into a tonic component (responsible for producing anti-gravitational joint torque for maintaining postural stability) and a phasic component (responsible for producing dynamic joint torque by including the effect of velocity and acceleration/deceleration of movement) allows for better evaluation of muscle activation and subsequently muscular synergies (7, 20, 21).

The present study, also the first to the best of our knowledge, examined the tonic and phasic components of trunk muscle synergies and their roles during flexural trunk movements. The aim was to test our hypothesis that the extracted time shifts following the phasic isolation of muscle activity can better explain the temporal patterns of synergistic activation.

## Materials and Methods

### Trunk and lumbar modeling

In this study, the trunk was modeled as a 3D inverted pendulum system integrated with a ball and socket joint at its base, located at the L5-S1 level, and controlled by 18 lumbar muscles (22, 23). A gravitational force of 350 N, concentrated at the center of mass, was applied at 35 cm above the pendulum’s joint (24). The following dynamic equation (Eq. 1) (25-27) describes trunk movement:

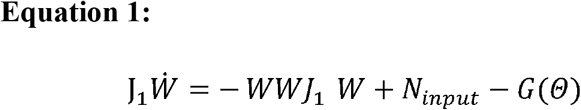

Θ is the vector of the trunk movement angle in the reference coordinates. W and *N*_*input*_, respectively, are angular velocity and muscular torque vectors in the coordinates connected to the body.*J*_1_ and *G*(*Θ*) are the inertial matrix and the gravitational torque vector of the trunk, WW are the Skew-symmetric matrix derived from the W gyration. *-WWJ*_1_*W* is the term of torque derived from the coriolis force. The maximum muscle velocity for muscles is *L*_0_/*τ*, which is the length of muscle relaxation and *τ* = 0.1 (sec). The radius of gyration is considered to be *R*_*x*_ = *R*_*y*_ = 0.44*m* and *R*_*z*_ = 0.1 m. *σ*_*max*_ = 70 *k Pa* is considered. The number of discretization points in this study is 60 points in the time interval of movement.

### Muscle modeling

The muscular forces involved in trunk movement were simulated using a three-element Hill muscle model. The 18 muscles included in this simulation were: internal oblique (IO), external oblique (EO), latissimus dorsi (LD), longissimus thoracic (LT), iliocostalis lumborum (IL), and rectus abdominus (RA), on both right and left sides of the body, respectively. The muscles and their physiological and geometrical properties, including lines of action and cross-sectional areas, were in line with previous studies (22, 28). Details of the modeling process and validity of the spine model were reported in our previous work (28).

### Computing optimal control and optimization

A minimum energy-jerk cost function (Eq. 2) was employed to solve the optimal control problem. Combining minimum jerk and energy in the cost function allows for time-optimized smoothed movement and provides an energy-efficient model (29).

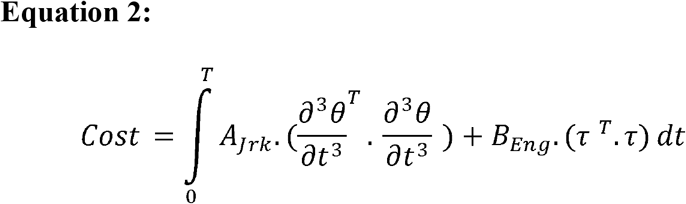

*τ* is the torque vector derived from muscle force around the joint. *A*_*jrk*_ and *B*_*Eng*_ are respectively the weighting coefficients of the minimum jerk and the minimum energy, both of which are equal to 0.5.

The numerical solution in this study was formulated based on the pseudo-spectral method (30). Position, velocity, and acceleration of the spine at the beginning and end of the movement path, as well as muscle activity in the range of 0 to 1, were considered as boundary conditions. Assuming no axial torsion, the torsional torque was neglected in the movement equations. The equations were then solved by trial and error with the goal of performing the movement in the shortest time possible with muscle activities below saturation levels. The SNOP solver in Tomlab PROPT (The Professional Optimal Control Software, MATLAB (Mathworks, Inc., Natick, MA, USA)) was employed (31). MATLAB’s “fmincon” function was also used to determine the level of muscle activity.

### Simulation and determination of boundary conditions

Twenty-four lumbar plane-bending movements (8 directions × 3 velocities) were modeled assuming no axial torsion occurring around the spine. The eight movement directions comprised of 4 main flexion/extension, right/left lateral bending, and 4 bending movements in the combined planes (Fig 1). These planes were placed in a 45-degree position relative to both the frontal and sagittal planes. In this study, movement in a specific direction at a certain velocity was defined as a “task”, where a total of 24 different tasks were modeled to investigate trunk muscle activation. Based on literature, 60, 45 and 22 degrees were estimated as ranges of motion (ROM) in flexion, right and left lateral bending, and extension, respectively (32). Using the curve fitting toolbox in MATLAB, the ROM was also estimated for four combined directions, 40 degrees for tasks 2 and 8 and 20 degrees for tasks 4 and 6 (Fig 1).

**Fig 1.**
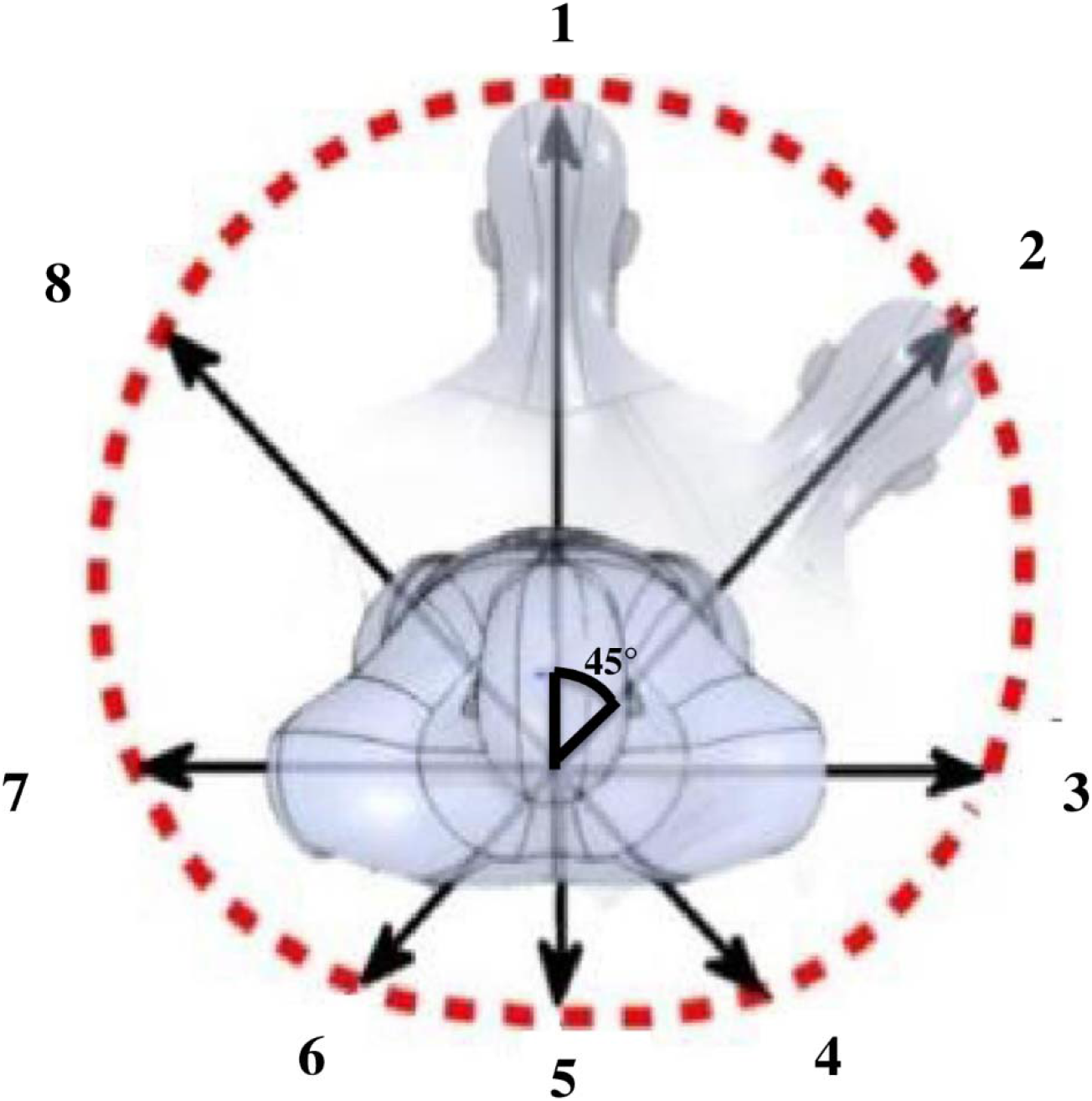
Movement planes modeled. Main planes: 1) 60° of flexion in the saggital plane, 3) 45° right bending, 5) 22° extension, 7) 45° left bending. Combined planes: 2) combination of flexion and 40° right bending, 4) combination of extension and 20° right bending, 6) combination of extension and 20° left bending, 8) combination of flexion and 40° left bending.

Three different path traversal times, including eight predefined directions, were modeled to evaluate the effect of velocity on muscle synergies (0.75, 1 and 2 seconds). Three levels of velocity were consequently obtained for each movement. Since the movements were performed in eight directions, a total of 24 movements were recorded in the model and muscle data was extracted. Initially, the body was assumed to be in an anatomical state of relaxed neutral standing, and bending was performed in a fixed direction during a constant time period. The velocity and acceleration at the initial and final stages of all movements were considered zero.

### Time-varying synergies

As previously mentioned, the muscle activity pattern was divided into two components: tonic (influenced by gravity) and phasic (proportional to acceleration/deceleration). By adopting this approach from d’Avella et al. (7), the time-varying synergies were calculated as follows by Eq. 3:

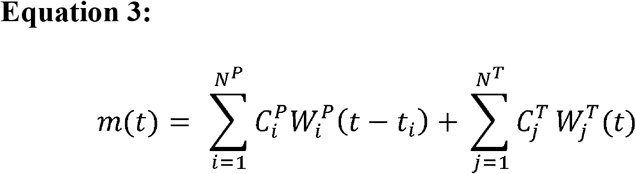

m (t) is the function of a muscle activity relative to time pattern. This function consists of two parts: the sum of phasic synergies with superscript “P” and tonic synergies with superscript “T”. The number of phasic synergies is i and the number of tonic synergies is j. As can be seen, the muscle pattern obtained from the phasic part can be transmitted by applying a time-shifting coefficient *t*_*i*_, during the time to find the best overlap with the pattern desired to perform the movement. It should be noted that the coefficients of phasic and tonic synergies 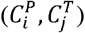 remain constant over time.

Initially, the muscle activation levels in the S^th^ movement were defined in a matrix in the form of M_*s* (*NM*×*K*)_, in which NM = 18 is the number of muscles and K = 60 is the number of discretization movement steps. S is the number of movement and can be varied from 1 to 24 (M = [M_1_,…,M_24_]).

Subsequently, each of the phasic and tonic synergies was defined as W_(NM×J)_ matrix, in which J is the number of discretization steps for synergies during the movement (assumed to be J = K = 60 in this study). Therefore, given the number of synergies for both phasic and tonic synergies of N, the total synergy matrix can be defined as W = [W_*P*1_,…, W_*PN*_, W_*T*1_,…, W_*TN*_].

Using the Kronecker Delta function, the “H” matrix, which contains the effects of both the amplitude and time delay, is defined by Eq. 4:

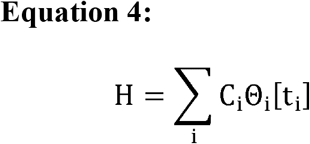

Where Θ_i_[t_i_] is the time-shifting matrix that contains 1 (for along a diagonal from start to finish i_th_ synergy) and 0. “i” is the number of synergy that vary from 1 to 4.

Hence, Eq. 3 was rewritten into a matrix form (i.e., Eq. 5). The squared reconstruction error between muscle activation and the reconstructed signal from extracted synergies is shown in Eq. 6.

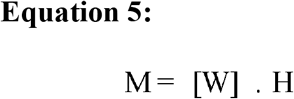

Where matrix M is muscle activation data matrix, W is muscle synergy matrix, and matrix is the function of amplitude coefficient and time delay for N synergies.

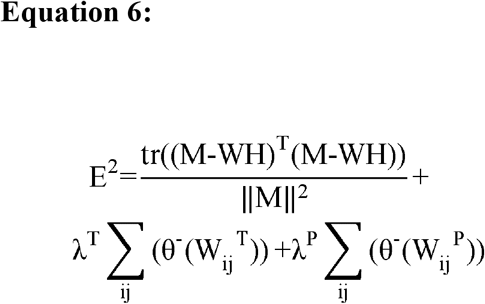

*E*^2^ is the squared reconstruction error equals the square of the difference in the regenerated muscle activation from the extracted synergies and original data, divided by the magnitude of the second power of the original data. Given the non-negative assumption of the value of synergies, two θ^-^(W_ij_)^*P*^ and θ^-^(W_ij_)^*T*^ functions are added to this equation to eliminate negative values in the W matrices, and λ^T^, λ^P^are the effect coefficients for increasing or decreasing the effect of these values on the main cost function.

The exact extraction procedure of synergies is presented as follows (7):

1. Detecting the synergy delays: Based on the maximization of the cross-correlation of the synergies with all episodes of data by using the matching pursuit method (33).
2. Updating the scaling coefficients: Given the non-negative phasic and tonic synergy coefficients, the scaling coefficients are calculated simultaneously by the non-negative least squares method of MATLAB software.
3. Updating synergies: Using the Steepest Descent optimization method and the error function (Eq. 6), the phasic and tonic synergies are updated simultaneously.

### Determining the suitable number of muscle synergies

We used *R*^2^ and mean squared error (MSE) versus N plot and the “knee-point” method (29) to determine the number of suitable synergies (N*) (Eqs. 7 and 8, respectively). N* was selected as the number for which the *R*^2^ and MSE curve has a considerable change in slope and where adding an additional synergy does not increase *R*^2^ by >5% (30-32).

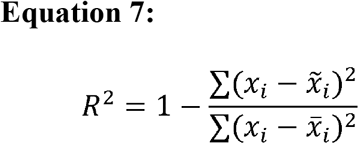

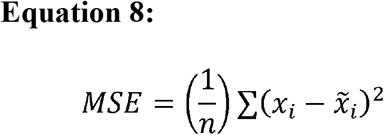

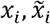 and 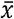 are initial data extracted from modeled movements, reconstructed data and mean value of *x*_*i*_, respectively. n is the number of the data sample.

## Results

Fig 2 shows the simulated kinematic profiles (position, velocity, and acceleration) of the trunk movement for 60° flexion from an upright posture (main plane no. 1 in Fig 1) during 1 second. The figure reveals a bell-shaped velocity curve and acceleration/deceleration patterns which reflect the co-activation of different muscle groups proportional to each part of the movement (Fig 2a). It also demonstrates that the flexor muscles are activated first to provide the initial acceleration required for the trunk (Fig 2b). When the muscles’ torque reaches zero, the extensor muscles are activated to oppose the movement ending at 60° flexion. In particular, the LT and IL muscles play the most significant roles in the deceleration of movement. The muscle activation profiles seen here are consistent with the results of Zeinali-Davarani et al.’s model on 60° trunk flexion during 1 second (22).

**Fig 2.**
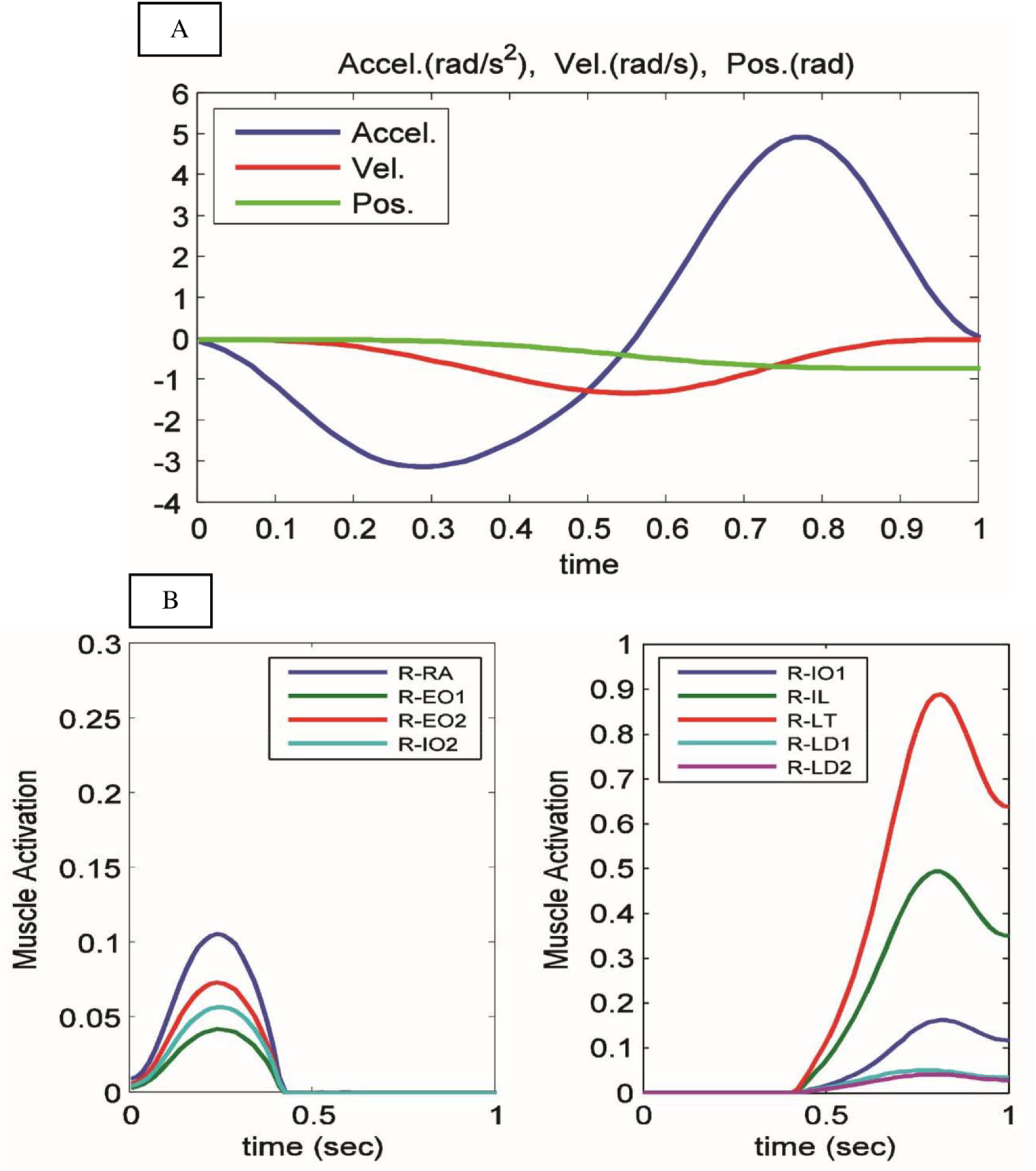
(A) An optimal movement path, velocity and acceleration of movement using the cost function, a combination of minimum energy and minimum jerk with equal coefficients. (B) The flexor and extensor muscle activity pattern of the right half of the body for the cost function. Given the symmetry of the model, only the right muscles of the body are shown.

According to Fig 3, reconstruction as a function of the number of synergies reached 75% by using 4 synergies based on the R^2^ criterion. There was a reduction in the diagram’s slope with this number of synergies. According to the MSE criterion, the error of reconstruction was below 2 ×10^−3^ at four synergies, and it was clear that increasing this number led to no significant changes at the error level. Therefore, four synergies were selected for each of the phasic and tonic synergies in this study.

**Fig 3.**
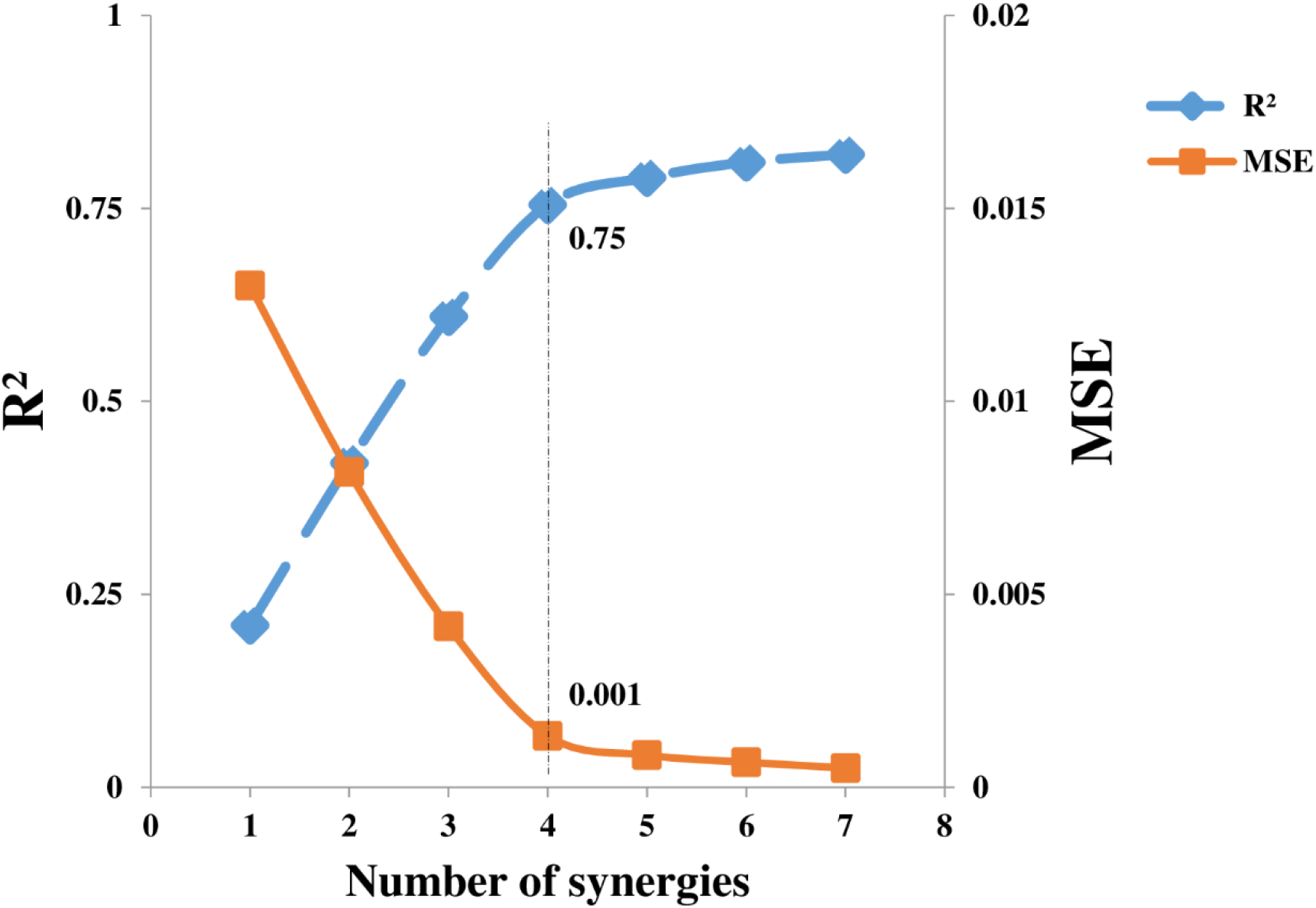
Reconstruction of primary data variation based on the number of synergies through two criteria of R^2^ (the blue line-axis of numbers on the left of the diagram) and MSE (the red line-axis of the numbers on the right side of the diagram). The pattern shows a change in slope at 4 synergies for both criteria.

The right and left abdominal flexor muscles reached peak activation in the first half of the movement in the first phasic synergy, while most back extensor muscles were activated and reached their peak in the second half of the movement (Fig 4). The trunk, therefore, accelerated towards flexion in the beginning of the movement, while the extensor muscles were activated in the second half of the movement during deceleration as synergy participated in the movement control. A similar acceleration scenario was observed in the second phasic synergy. However, here only the right abdominal and left back muscles were activated, suggesting that bending to the right side has led to the activation of the agonist muscles and the muscles in the opposite direction of the antagonists at the beginning and end of the movement, respectively. In contrast to the first synergy, the extensor muscles were activated first in the third synergy, possibly to move towards extension from a static state. The peak muscle activity range in this synergy was smaller than the first synergy, probably because the range of motion modeled for the extensor tasks was less than the flexural tasks (for instance, 22° for task 5 vs. 65° for task 1). Furthermore, a relationship was observed between the level of activity and the bending amount. The timing in the muscle co-contraction in synergy 4 seemed to be the opposite to that of synergy 2, suggesting contribution to bending on the left side.

**Fig 4.**
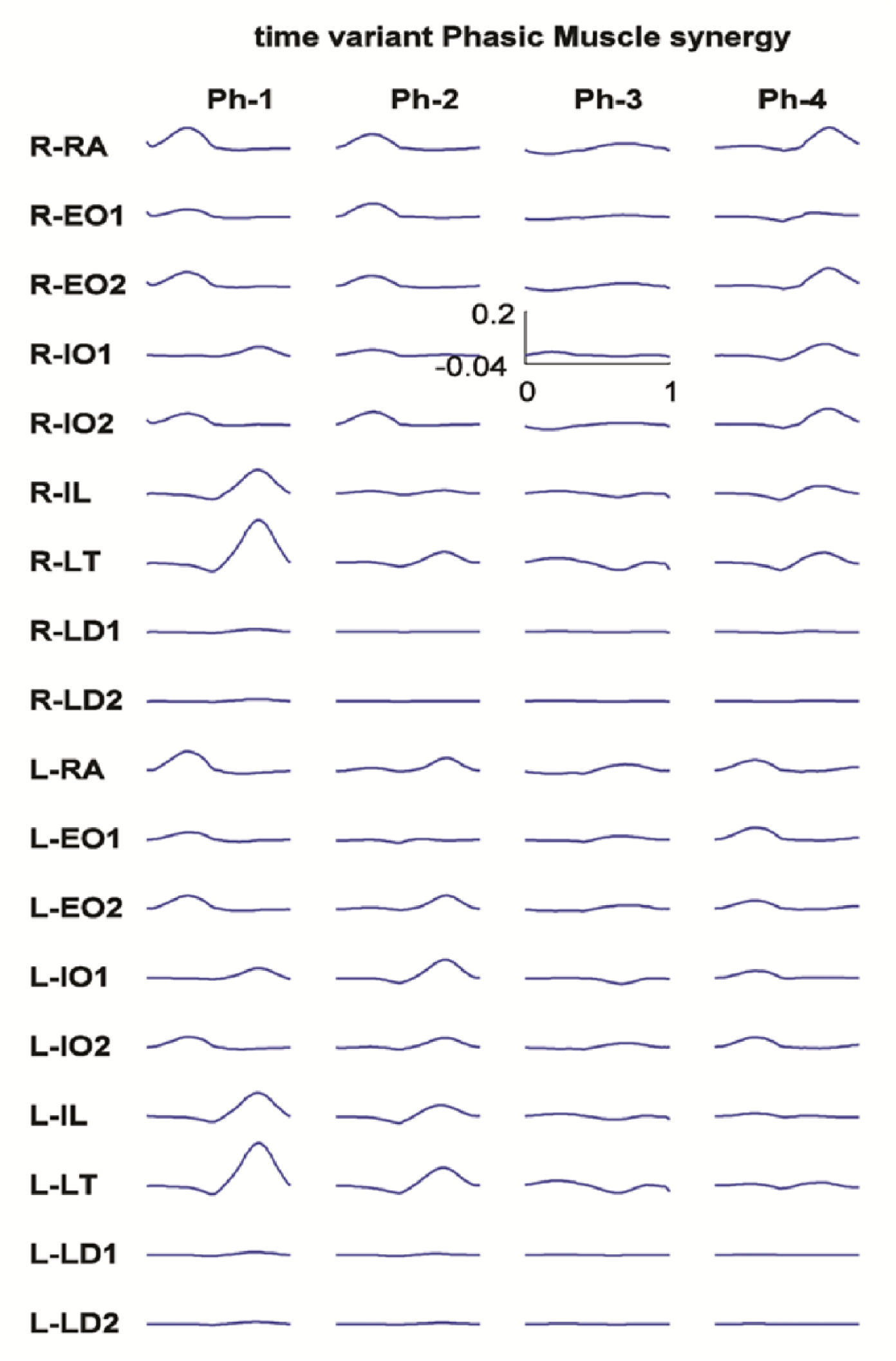
Muscle activation pattern in four-time variant phasic synergies for 18 muscles

The present study revealed that synergy 1 is predominantly active in forward flexion, while synergy 3 plays a major role in backward extension (Fig 5A). It is worthwhile noting that the predominant synergy in each direction was the synergy with the largest amplitude coefficients in these directions. The synergy time shifts are demonstrated in Fig 5B. The figure depicts the different time shifts in different directions. For example, the value of the time shift of synergy 3 is close to zero in the predominant direction (extension), while it is close to -1 in the non-dominant direction (flexion). A time delay which is estimated at zero for a particular synergy implies that the synergy activates at the beginning of the movement, while a value reaching -1 suggests earlier activation. Therefore, when synergy 3 has zero shift in the dominant direction reflects that all of the synergy participates in the movement, in line with the dominant nature of the synergy in this direction. Fig 6 illustrates the effect of amplitude coefficients and time shift on the reconstruction of a muscle activity pattern by four desired synergies as an example.

**Fig 5.**
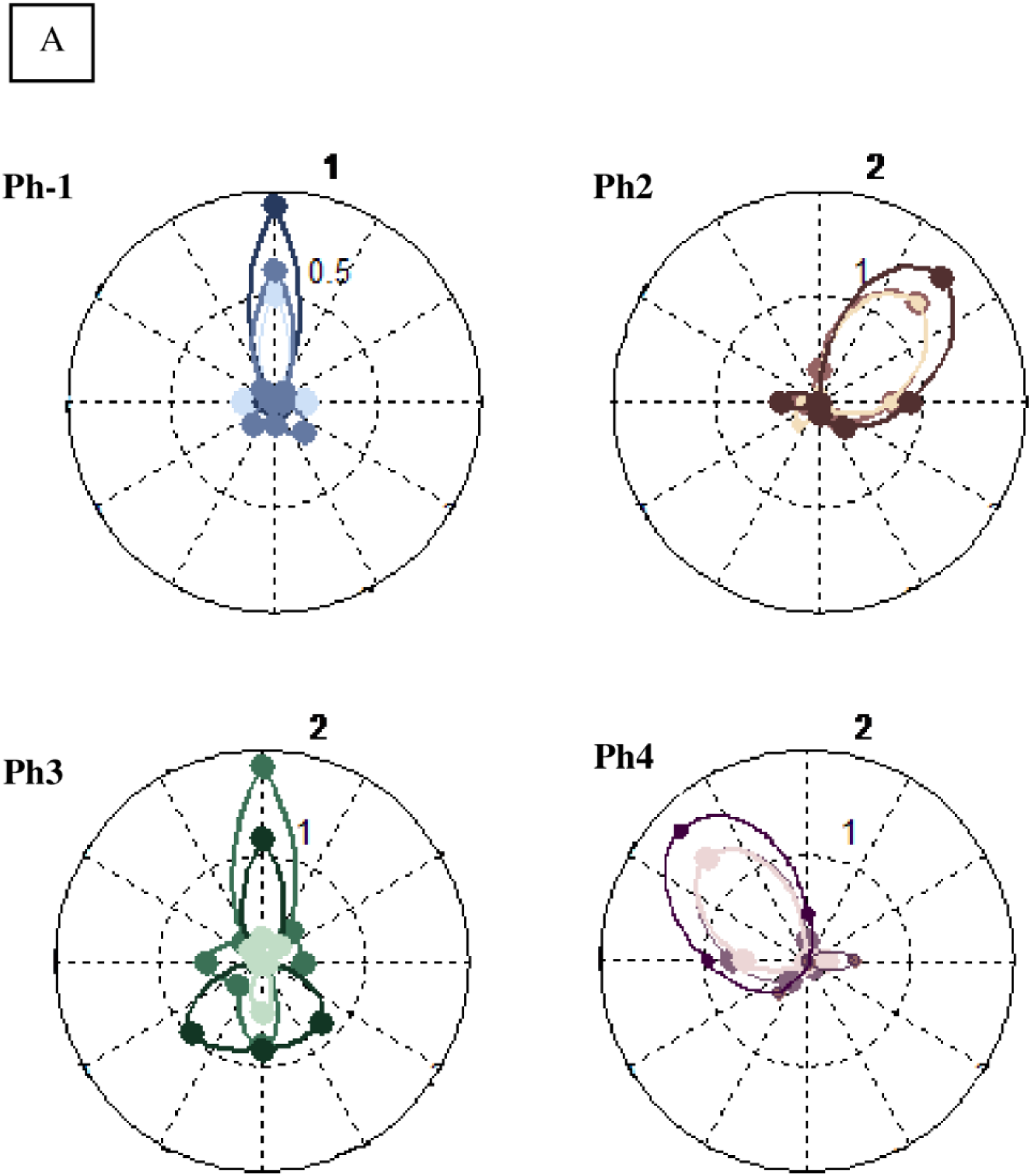
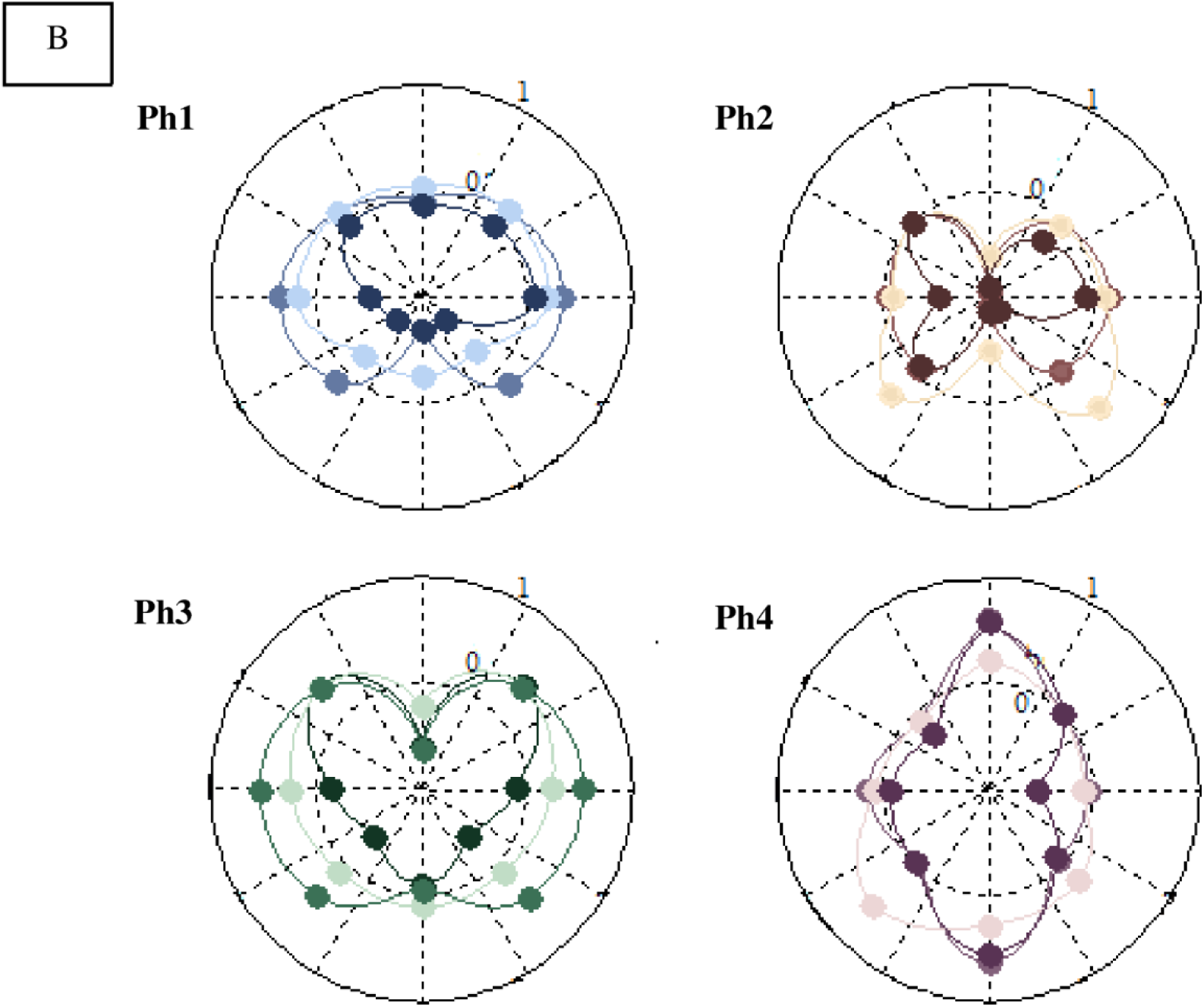
(A) Amplitude coefficients of four phasic synergies in polar diagram for flexion-extension in eight different directions and three different velocities modeled. The synergies one to four are shown with colors blue, brown, green, and purple, respectively. Larger motion velocities are displayed in darker colors. (B) Time shifts of four phasic synergies in a polar diagram for flexion-extension in eight different directions and three different velocities modeled. Synergies one-four are shown in blue, brown, green, and purple colors, respectively. Higher motion velocities are displayed in darker colors.

**Fig 6.**
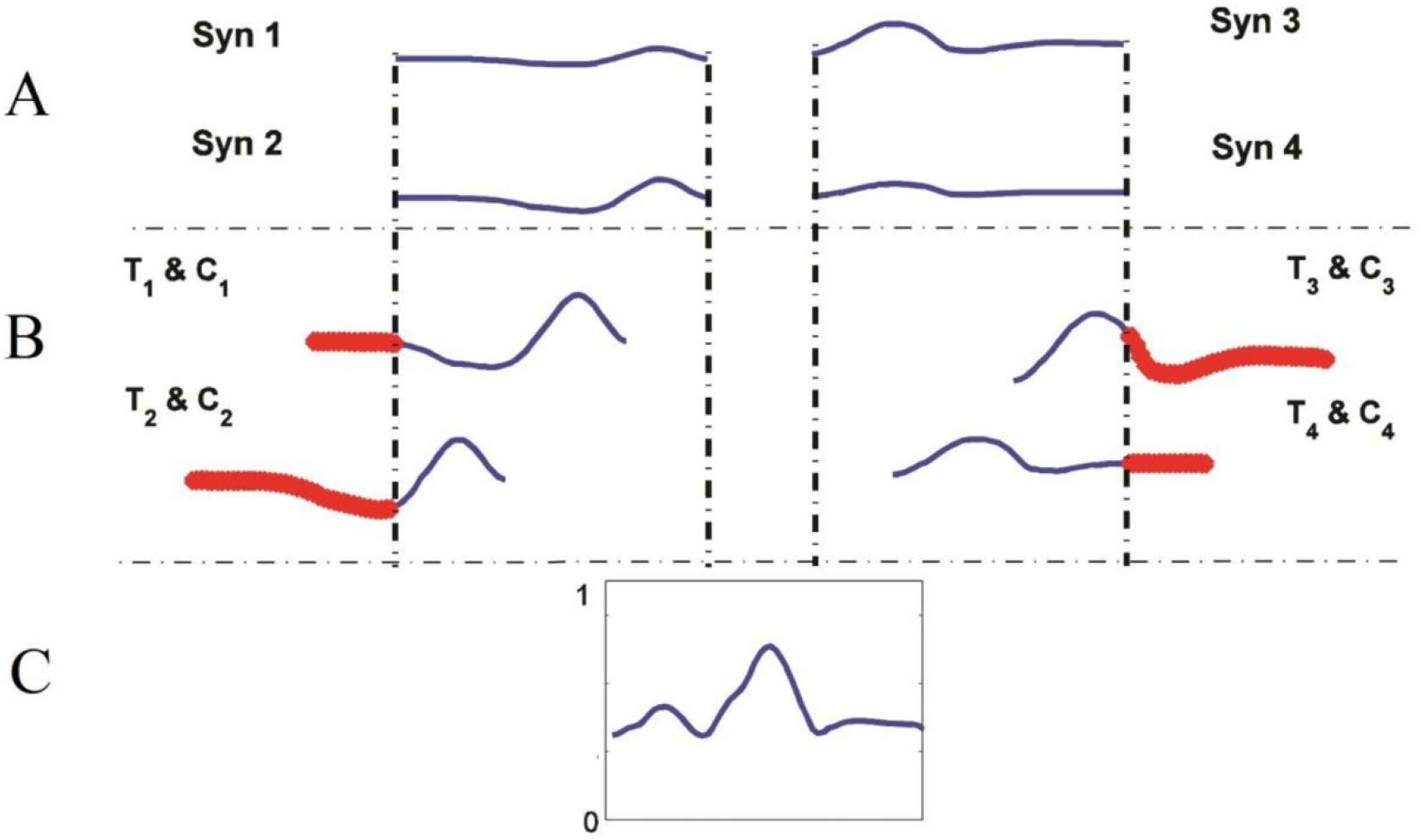
A schematic of the effect of time shift and amplitude coefficients on phasic synergies for reconstruction of a desired muscle activity. (A) Four time-variant synergies were selected with the desired duration of activity. (B) Each synergy was scaled and shifted with a specific amplitude coefficient (*C*_*i*_) and time shift (T_i_). (C) Reconstructed activity of a desired muscle with linear combination of the modeled scaled and shifted synergies.

Tonic synergies have much larger amplitude coefficients as compared to phasic synergies due to the weight of the trunk (Fig 7). However, similar to the phasic synergies, each tonic synergy has certain groups of active muscles. For example, the activities of the lumbar muscles in the first synergy and the abdominal muscles in the fourth synergy increased over time (Fig 7).

**Fig 7.**
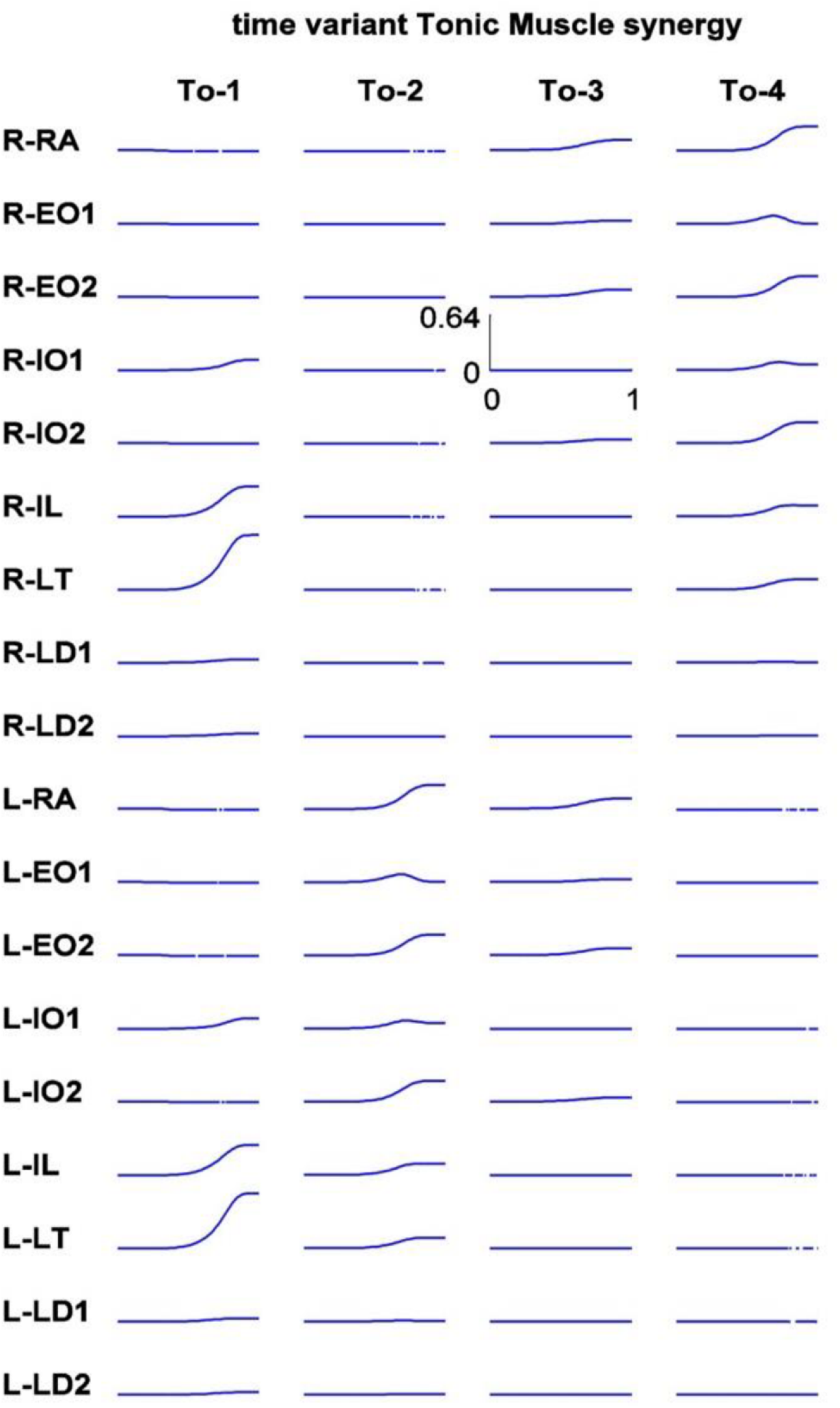
Four-time variant tonic synergies for 18 muscles

To understand the participation of tonic synergies in different movements, the magnitude of the coefficient in each task should be observed. Examining the amplitude coefficients of tonic synergies (Fig 8) reveals that each synergy is dominant in a part of the space, and that the coefficients are completely modulated with the direction of movement (modeled task). However, speed does not have a clear impact on these coefficients, and it is not possible to determine exactly how the changes in speed have caused the changes in the coefficients of tonic synergies.

**Fig 8.**
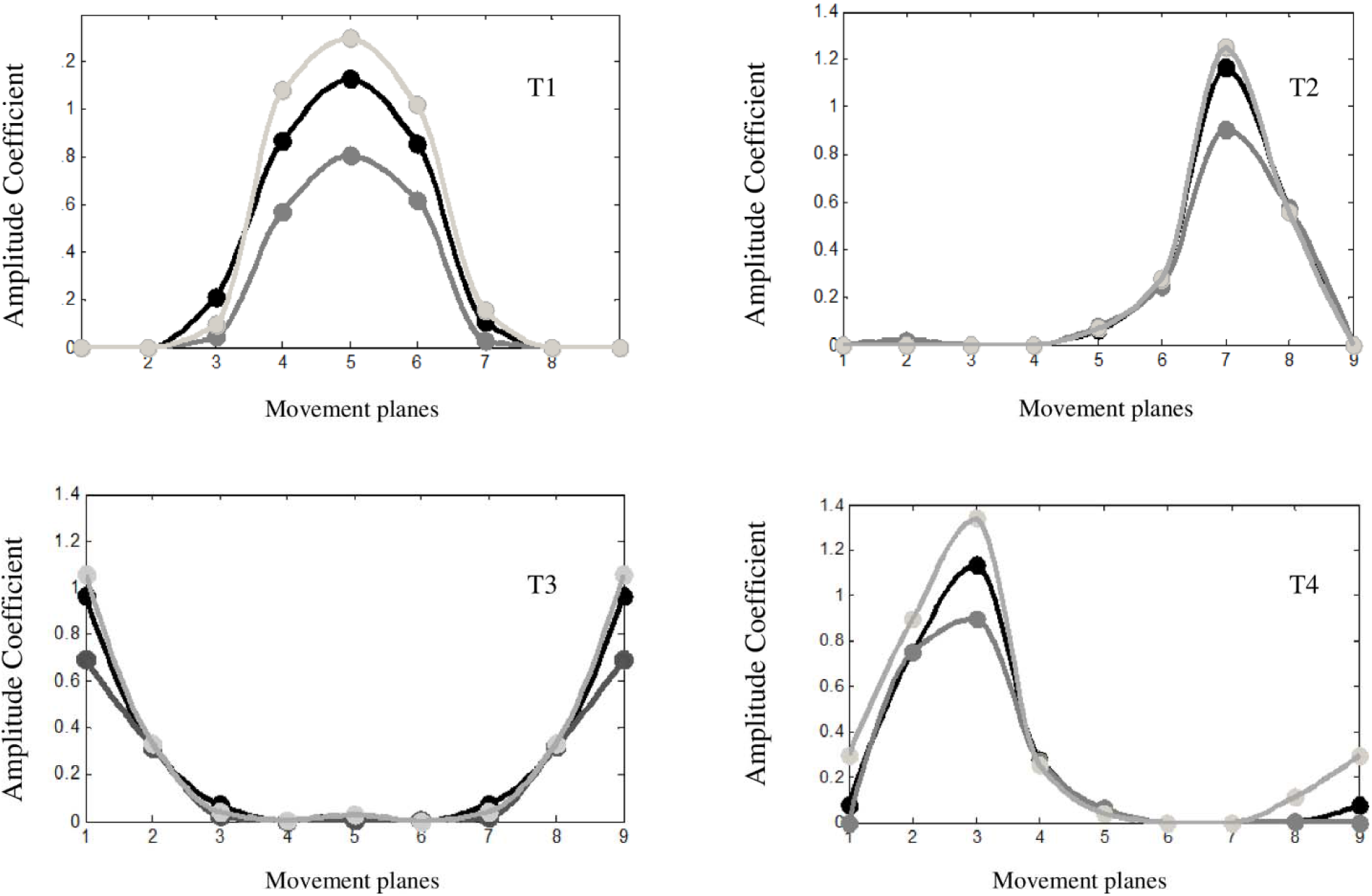
Amplitude coefficient for 4 time-varying tonic synergies (T-1 to T-4) in various directions and velocities. Higher motion velocities are displayed in darker colors. The eight values of the coefficients for each synergy (colored dots) are connected by a periodic cubic spline interpolation curve. Modeled movements planes are named 1 to 8 and the number 9 is as same as number 1 to crate symmetry at the beginning and end of the patterns

Fig 9 depicts the different coefficients of the tonic synergies in a polar diagram at a specific speed (performing each motion in 1 second). As can be seen in the figure, synergies 1 to 4 are predominantly activated in the range of space related to forward flexion, right lateral flexion, extension, and left lateral flexion, respectively. In combined planes, the contribution of two or more synergies is needed to create a movement.

**Fig 9.**
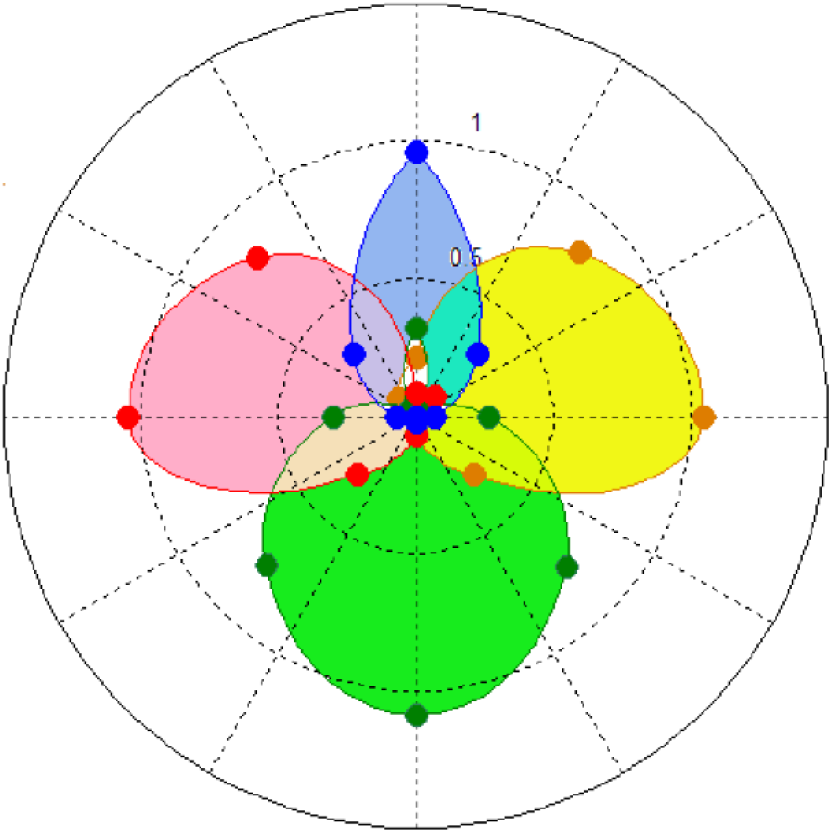
Amplitude coefficients of four tonic synergies in pole diagram to create a motion at the highest velocity (performing each motion in 1second) in eight directions modulated; synergies one to four are shown in blue, yellow, green and pink colors, respectively.

To show the impact of synergies on muscle data reconstruction, a task other than the 24 initial tasks was modeled, and the activities of all 18 muscles were obtained. The task included a 45-degree bending to the right in a 30-degree plane relative to the sagittal plane, which was placed between directions one and two (Fig 1). The task was modeled for a 1 second duration of movement. The second step involved reconstructing the activity of muscles using synergies, where the first and second tonic and phasic synergies active in this area of space were selected for reconstruction. The amplitude coefficients for the first and second phasic synergies were considered 0.1 and 0.5, respectively, whereas a value of 0.5 was used for both the first and second tonic synergies. While the time shift was considered 0 for the first phasic synergy, the second synergy was activated 0.1 second earlier. On the other hand, no time shift was considered for tonic synergies (Fig 10). The R^2^ criterion confirmed that more than 75% similarity between the main and reconstructed data.

**Fig 10.**
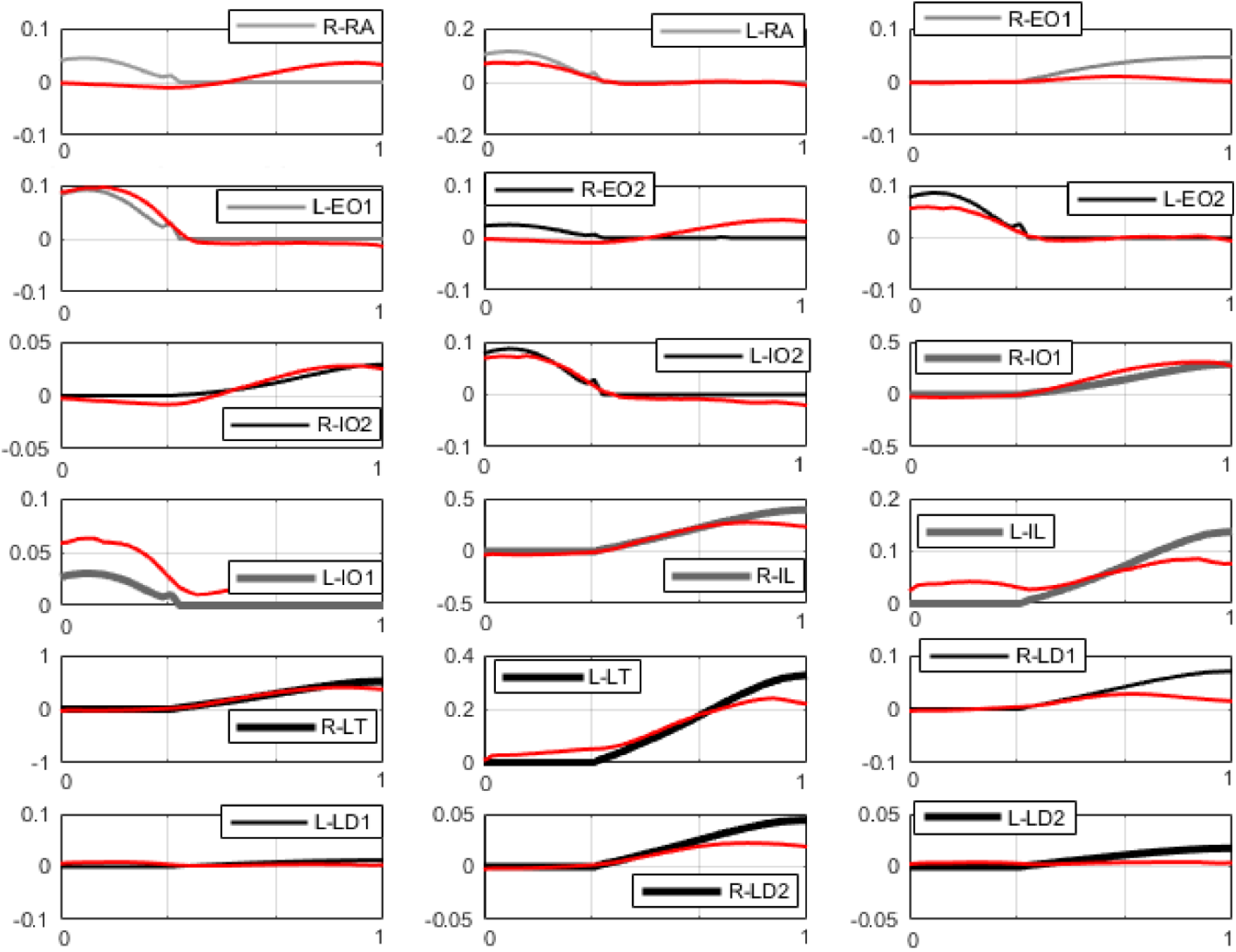
Data reconstructed by time-variant synergies compared to the data obtained from the model for 18 muscles involved in 45° right bending in a 30° plane relative to the sagittal plane. The reconstruction data and the main data obtained from the model are displayed with red and dark lines, respectively. Using R^2^, the performance of reconstruction was higher than 70%.

## Discussion

The main goal of this work was to bridge the gap in literature regarding the impact of velocity and direction of motion of the trunk on amplitude coefficients and the onset time of trunk muscle synergies. The synergies were further separated into tonic and phasic components for better understanding of their relative contribution and respective roles in the control of flexural trunk movements. More specifically, the time-varying synergies of 18 simulated lumbar muscles during spinal flexion-extension movements with different velocities and directions were obtained, and an optimization model using a combined minimum energy-jerk cost function was formulated.

Overall, our results suggest that it is conceivable for the CNS to employ a series of organized, modulated, time-variant and amplitude-varying synergies as a series of fundamental functional units to control complex motions, such as lumbar flexion-extension in different directions and at various speeds. In our optimal control model, the Bang-Bang control of the transmitted signals was able simulate a series of neural stimuli and activate the muscles according to the prescribed cost function, which ultimately resulted in muscle activity and acceleration-deceleration of movement as model outputs (Fig 2A and Fig 2B). The biphasic and triphasic shapes of the muscle activity patterns (Fig 4), revealed that the control signals reach the muscles at different times of movement, and hence different muscle groups are activated with various delays. This is in alignment with the intermittent optimal control-based model developed by Leib et al. (29) who showed that the co-activation of agnostic and antagonistic muscles during different movement phases was proportional to acceleration/deceleration. Moreover, this form of activation (biphasic or triphasic) allows the synergy to be more involved in motion reconstruction, which may explain the parsimony and low-dimensionally nature of time-variant synergies, as compared to synchronous synergies (7).

To overcome the high torque caused by the trunk’s weight during bending, the muscles that were activated in the form of tonic synergies needed to increase their activation level. This was demonstrated in our model, where the level of muscle activities in tonic synergies (Fig 7) was higher as compared to the activation levels in the phasic synergies (Fig 4) (0 to 0.64 vs. -0.04 to 0.2, respectively). Tonic synergies usually peak at the end of the movement when maximum trunk torque is applied to the lumbar region. However, due to acceleration-deceleration, phasic synergies were more active during the movement. Our results confirm that tonic synergies are generally monophasic, and hence considering time shifts does not help to understand them better. In the present study, a dominant synergy was observed in the main directions, but in the combined planes, two or more synergies cooperated with each other (Fig. 5A and Fig. 9). As per our hypothesis, the amplitude coefficients of the phasic and tonic synergies depended on both the direction and velocity of movement. The increase in velocity only led to an increase in the coefficients of phasic synergies (Fig. 5A), but not in tonic synergy (Fig. 8). Increased coefficients of phasic synergies can be due to the need for higher muscle stiffness at higher velocities. This finding was also reported in the study of d’Avella et al. regarding the phasic synergies of the shoulder muscles, where the increase in velocity was accompanied by an increase in the amplitude coefficients (16).

The time shifts obtained in this study were sensitive to the direction of motion, where each synergy was activated earlier or later in part of the movement. The time shifts in the dominant synergy were close to zero (Fig. 5B). The results also demonstrated that the higher the velocity, the faster the activation of the phasic synergy (Fig. 5B). This is probably due to increasing the velocity which requires increased muscle activity, especially at the beginning of the movement (acceleration phase). In agreement with literature, the agonistic synergies were activated sooner to produce faster movements (34).

Some limitations of this study should be delineated here. This work did not incorporate the effect of joint stability, which should be considered in future research as a constraint to our model to examine how muscled cooperate to maintain stability. The results of the present study are based on modeling with all its advantages and inherent limitations. It is worth noting here that for a more comprehensive analysis, further biomechanical modeling considerations should be taken into account. These include intra-abdominal pressure, multi-segment modeling of the spine, inclusion of passive tissues, inclusion of muscle via points, more detailed musculotendon modeling, etc. This study can also be improved by using EMG data to study the time-varying synergies of lumbar muscles during functional tasks.

## Conclusion

Our findings suggest the plausibility of a time-variant muscle synergies control scheme by the CNS for optimal trunk movement during flexion-extension in 3D at different speeds. The separation of muscle activities into two tonic and phasic components provides a more realistic assessment of the impact of kinematic (velocity and acceleration) changes on movement dynamics and control. The results of this work can be applied to other fields such as robotics, control of smart prostheses, biomechanics, and rehabilitation. Importantly, this study confirms that the time-varying synergies approach may shed more light on the control strategies of the trunk neuromuscular system and provide insight into the mechanisms underlying locomotor robustness and flexibility.

## Acknowledgment

The authors are grateful for the invaluable assistance of Dr. Bahman Nasseroleslami in preparing this paper.

